# Generalized differential equation compartmental models of infectious disease transmission

**DOI:** 10.1101/777250

**Authors:** Scott Greenhalgh, Carly Rozins

## Abstract

For decades, mathematical models of disease transmission have provided researchers and public health officials with critical insights into the progression, control, and prevention of disease spread. Of these models, one of most fundamental is the SIR differential equation model. However, this ubiquitous model has one significant and rarely acknowledged shortcoming: it is unable to account for a disease’s true infectious period distribution. As the misspecification of such a biological characteristic is known to significantly affect model behavior, there is a need to develop new modeling approaches that capture such information. Therefore, we illustrate an innovative take on compartmental models, derived from their general formulation as systems of nonlinear Volterra integral equations, to capture a broader range of infectious period distributions, yet maintain the desirable formulation as systems of differential equations. Our work illustrates a compartmental model that captures any Erlang distributed duration of infection with only 3 differential equations, instead of the typical inflated model sizes required by differential equation compartmental models, and a compartmental model that capture any mean, standard deviation, skewness, and kurtosis of an infectious period distribution with merely 4 differential equations. The significance of our work is that it opens up a new class of easy-to-use compartmental models to predict disease outbreaks that does not require a complete overhaul of existing theory, and thus provides a starting point for multiple research avenues of investigation under the contexts of mathematics, public health, and evolutionary biology.

## 1. Introduction

The compartmental model of Kermack and McKendrick [1–3] is arguably one of the greatest development in disease modeling. The formulation of this model, in its original form as a system of nonlinear Volterra integral equations [4], provides a general characterization of the transmission cycle between susceptible individuals and a disease that propagates throughout an environment [5]. Despite this generality, the vast majority of disease modellers prefer differential equation compartmental models [6]. While this particular formulation of compartmental models has distinct advantages, such as the non-requirement of specialist knowledge to implement and well-developed numerical methods for their computation [7], they are in fact a special case of the aforementioned system of nonlinear Volterra integral equations [8]. Specifically, one obtains differential equation compartmental models from the nonlinear Volterra integral equations by imposing only two assumptions: 1) the infectiousness of a disease corresponds to disease incidence, and 2) the duration of infection follows either an exponential or an Erlang distribution. Unfortunately, the vast majority of diseases do not have a duration of infection that follow these distributions [9–14], and force fitting such a distributional structure is known to have a massive effect on the behavior and quality of model predictions [15]. Regardless of these issues, the compartmental models obtained by the two traditional assumptions have undergone many extensions. A few noteworthy examples include modifying the force of infection to account for the saturation of infection in a population [16,17], behavioral characteristics [18–21], modification of the recovery rate to capture disease burnout [22], and the inclusion of additional disease stages [8]. Furthermore, the applications of these models have also grown considerably from just predicting a disease’s trajectory. Today, these traditional compartmental models are used to evaluate the health benefits and cost-effectiveness of public health policies and disease interventions [23,24], gauge the potential for disease virulence evolution [25], predict dominant influenza strains [26], investigate the complexities of disease co-infection [27], among many others. However, despite this growth in theory and application, the generalization of the very foundational assumptions that simplifies systems of nonlinear Volterra integral equations to differential equation compartmental models remains largely undeveloped.

In what follows, we propose new assumptions to simplify systems of nonlinear Volterra integral equations to systems of differential equations. The biological motivation for these new assumptions stem from the idea that a disease’s duration of infection distribution changes throughout an epidemic, whereas the infectious period distribution of a disease remains invariant. Consequently, we extend current models from solely tracking disease incidence to tracking the number of person-days of infected individuals. The motivation for the units of person-days is due to its use as a measurement in epidemiology for quantifying actual time-at-risk, and is calculated based the number of people and total time contributed. To account for tracking person-days, we make the assumptions in our analysis that 1) the total infectiousness of a disease corresponds to the product of disease incidence with a time-varying average duration of infection and 2) the duration of infection is distributed according to a non-homogeneous analog to the exponential distribution. Under these assumptions, we derive a novel class of differential equation compartmental models, provide model equilibria, and disease reproductive numbers [28].

Essential in the development of this new class of models is the use of survival analysis. Specifically the development of our novel class of models requires the hazard function and the mean residual waiting-time of a distribution [29,30], which is used to describe the time-varying average duration of infection. Therefore, we briefly outline some of the fundamental properties of these functions, in addition to their relationship to one another. We then demonstrate the consistency of our new class of models to traditional models when the duration of infection follows an exponential distribution. Next, we consider our model with a duration of infection that follows an Erlang distribution. We illustrate how this choice of distribution has a representation as either an ODE system of 3 equations, regardless of the Erlang distribution parameters, or as an ODE system that features a chain of infected individual equations, as is typical from the linear chain trick [9,31,32]. Finally, we consider a duration of infection that is Pearson distributed. In choosing the Pearson distribution, we develop a model that is capable of accounting for any possible mean, standard deviation, skewness, and kurtosis of the duration of infection. Thereby, we provide a simple approach for measuring how altering the infectivity profile of a disease, as described by the first four statistical moments, influences a diseases trajectory.

## 2. Methods

In what follows, we develop a novel class of ODE compartmental models to describe the progression of a disease throughout a population. To obtain such models, we impose new assumptions on the notion of infectivity used in the integral equations of Kermack and McKendrick. Before reducing the integral equations of Kermack and McKendrick to their differential equation counterparts, we briefly highlight the formal relationship in our assumptions in the context of survival analysis. Finally, we present the class of novel compartmental models, their equilibria, and basic reproductive number.

### 2.1 Traditional compartmental models

To begin, considered the general compartmental model of Kermack and McKendrick [1–3,5]. For this general compartmental model, we consider the number of susceptible individuals to be denoted as *S*, and the total infectivity of the disease (at time *t*) to be denoted as *ϕ*(*t*). We define *ϕ*(*t*) as the sum of the products of the number of individuals at a particular age of infection that remain infectious at time *t*, with the mean infectivity for that particular infection age. We also define *ϕ*_0_ as the total infectivity of the individuals initial infected with the disease at the start of the epidemic. In addition, we characterize the fraction of infected individuals remaining infected at time *t*, who where initially infected at time *x*, with the duration of infection distribution, as described by the survival function *P*(*t, x*). Furthermore, we also define the mean infectivity of an infected individual *t* − *x* units of time after the start of the infection as *π*(*t* − *x*), where 0 ≤ *π*(*t* − *x*) ≤ 1, and the mean infectivity of individuals in the population at time *t* who where initially infected at time *x* as

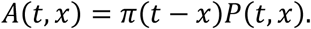

For simplicity, we assume a constant mean infectivity, *π*(*t* − *x*) = 1. Given that transmission is a function of both shedding rate and behaviour, it would be difficult to quantify changes in mean infectivity throughout an infection. Therefore, the progression of an epidemic throughout a population is described with the integral equations,

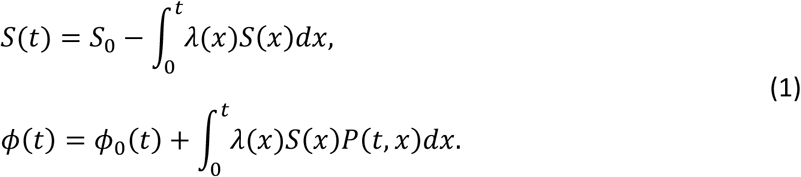

Here *λ* is the force of infection, which we assume to be

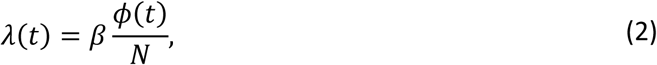

where *β* is the average number of contacts individuals in a population make per unit of time and *S*_0_ is the initial number of susceptible individuals at the start of the epidemic.

Traditionally, to reduce (1) to a system of differential equations requires that 1) the duration of infection follows the exponential distribution,

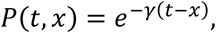

where *γ* is the recovery rate, and 2) that the infectiousness of a disease corresponds directly to the number of infected individuals, *ϕ* = *I*. Combining these assumptions, along with an additional compartment to track recovered individuals, *R*, transforms system (1) into

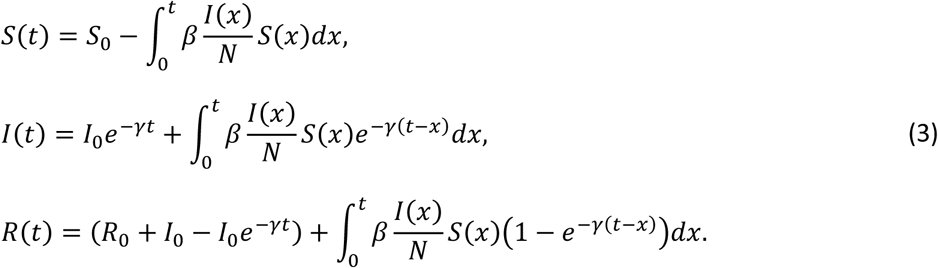

Here *I*_0_ and *R*_0_ represent the total number of infected individuals and removed individuals at the start of the epidemic. An important feature of system (3) is that it conserves the total population:

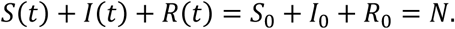

Differentiating system (3) with respect to *t*, and substituting the integral equation for *I*(*t*) for the remaining integrals yields the classic SIR system:

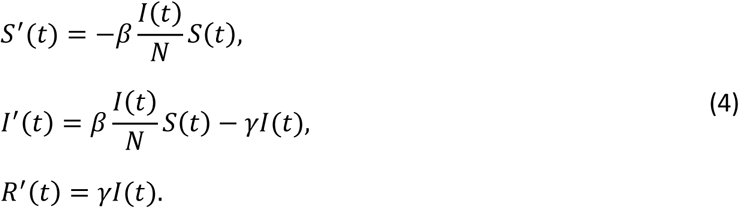

Alternatively, (1) reduces to a system of differential equations when 1) the duration of infection is the survival function of the Erlang distribution,

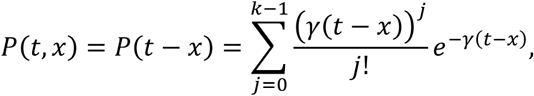

where *k* is a shape parameter that determines the total number of infection stages, 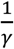 is the average duration spent in each stage, and 2) the total infectivity of the disease corresponds to *k* identical stages (in terms of the average duration spent in each stage) of infected individuals,

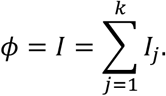

Combining these assumptions, along with an additional compartment to track recovered individuals, *R*, transforms system (1) into

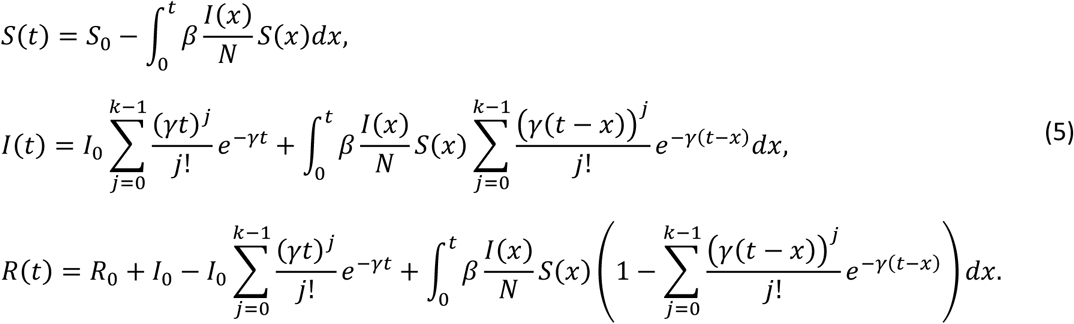

Equivalently, if the linear chain trick is applied, we have that

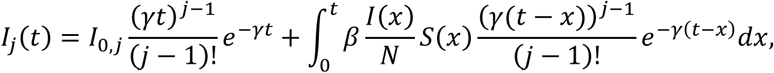

where *I*(*t*) = ∑_*j*_ *I*_*j*_(*t*).

An important feature of system (5) is that it conserves the total population:

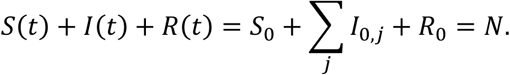

Differentiating system (5) with respect to *t*, applying the ‘linear chain trick’, and substituting *γI*_*j*_(*t*) as needed yields the classic SI^k^R system:

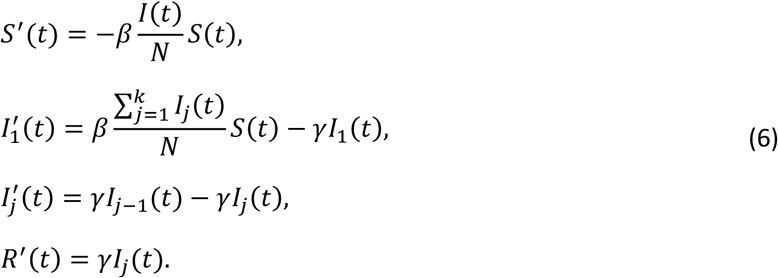

To obtain our novel differential equation compartmental models, we generalize the assumptions used to formulate system (5). We first assume the infectiousness of a disease corresponds to the product of the number of infected individuals with their average duration of infectiousness at time *t*:

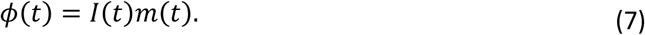

Here *m*(*t*) is a mean residual waiting-time [30]. Motivation for choosing *m*(*t*) over the typical (constant) average waiting-time stems from the fact that the composition of infected individuals *I*(*t*) includes individuals from different initial infectiontimes, and that individuals infected at different times are not likely to remain infectious for the same time period. By including *m*(*t*) in (7), our notion of infectivity is able to differentiatie between similar quantities of infected individuals that may (or may not) contribute differently to the spread of an epidemic.

In addition to assumption (7), we also assume that the duration of infection distribution corresponds to a non-homogeneous analog of the exponential distribution, namely the survival function given by

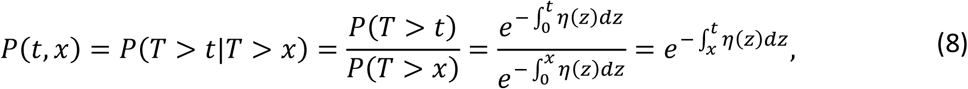

where *t* ≥ *x*, and *T* is a random variable that denotes the sojourn time in the infectious state.

### 2.2 The hazard function and mean residual waiting-time

Due to the importance of assumptions (7) and (8), we now provide a brief overview of the relationship between the hazard function, the mean residual waiting-time, and the survival function [29]. Consider a random variable *T* characterized by the survival function (8), which represents the probability of remaining infectious *t* units after becoming infectious at time *x*. The associated hazard function to *T* is given by

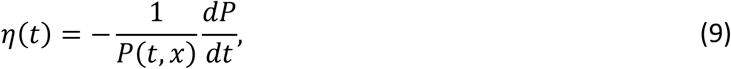

where *P*(*t, x*) is the survival function (for remaining infected) for individuals initially infected at time *x*.

Similarly, the mean residual waiting-time associated with *T* is also determined from (8) [29], and defined as

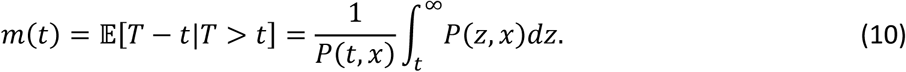

From the mean residual waiting-time (10), it is also possible to uniquely determine the hazard function (6) through the relation [29],

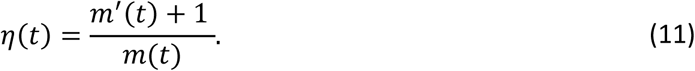

An important feature of the mean residual waiting-time (10) is that it is initially equivalent to the average duration of infectiousness, as

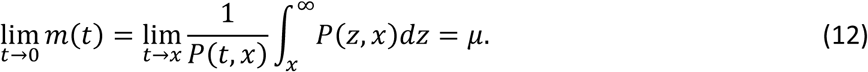

It follows from (10) and (11), and through the application of l’Hôpitials rule, that

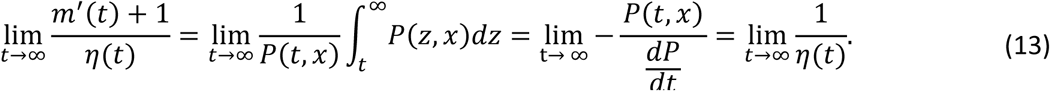

This implies if *η*(*t*) > 0 ∀*t* that

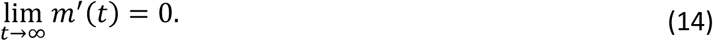

In addition to determining *η*(*t*) from *m*(*t*), the receprical relation is also possible. For convenience we restrict (8) to the family of distributions [29] that satisfy

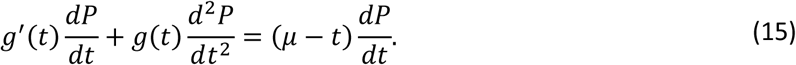

Note condition (15) ensures that both *m*(*t*) and *η*(*t*) exist and are finite [29]. Integrating (15) over [*t*, ∞), and using the property that 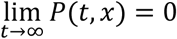, we obtain

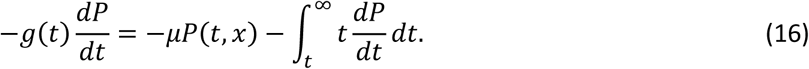

Noting that 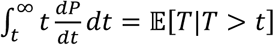, definitions (6) and (7), and dividing through by *P*(*t, x*) we obtain

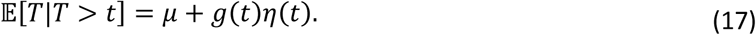

Thus, subtracting *t* from both sides we have that

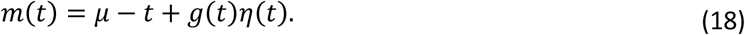

### 2.3 Novel compartmental models

Before presenting the novel compartmental model we briefly contextualize its differences in comparison to traditional compartmental models. The traditional compartmental models of (4) and (6) track the changes in the number of individuals suscpetible, infected, and recovered from disease, for a disease circulating in a population. The premise of our compartmental model is to instead consider tracking the change in the number of person-days. Specifically, we considers the individuals infected with a disease multipled by their average duration of time spent in the infected state. We also consider a similar setup for individuals suscpeptible to and recovered from disease. Therefore, to appropriately account for the temporal aspect of person-days we incorporate *m*(*t*), the mean residual waiting-time [29] into each compartment. We have that the person-days of the susceptible, infected, and recovered individuals in a population are governed by,

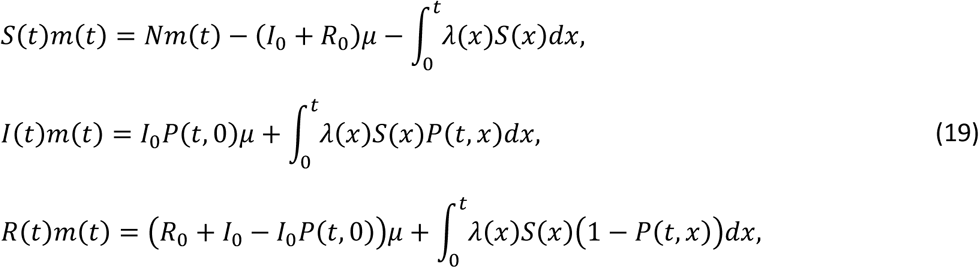

where *m*(0) = *μ* by (9), and *P*(*t, x*) is given by (5). Note, the term *Nm*(*t*) − (*I*_0_ + *R*_0_)*μ* accounts for changes in the time-varying reference frame for the total person-days of those susceptible to infection in the population.

Adding the equations of system (19) together, it follows that

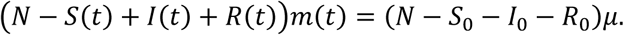

Thus, given *N* = *S*_0_ + *I*_0_ + *R*_0_ and *m*(*t*) ≠ 0 for all *t*, we have that

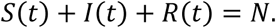

Imposing (7-10), taking the time derivative of system (19), and applying Leibniz rule for the derivatives of integrals as needed, we obtain

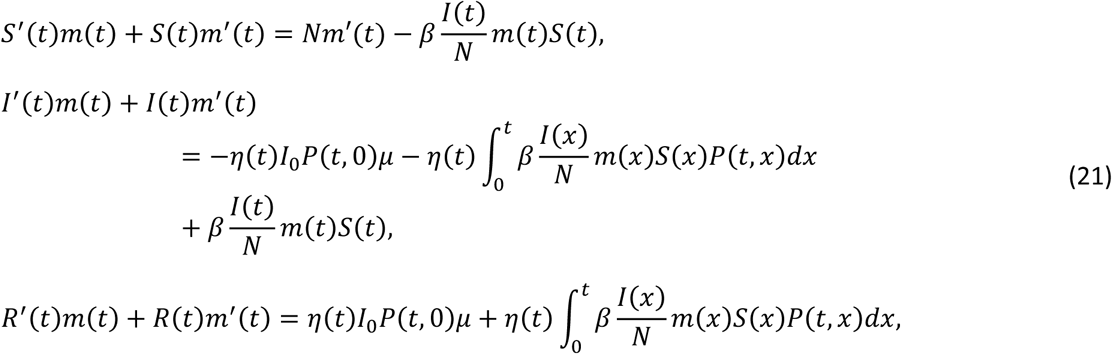

where *P*(*t, x*) is given by (8), and *η*(*t*) is given by (11).

Substituting (19) into (1) to eliminate the integrals, and isolating for each of *S*′(*t*), *I*′(*t*), and *R*′(*t*), we obtain:

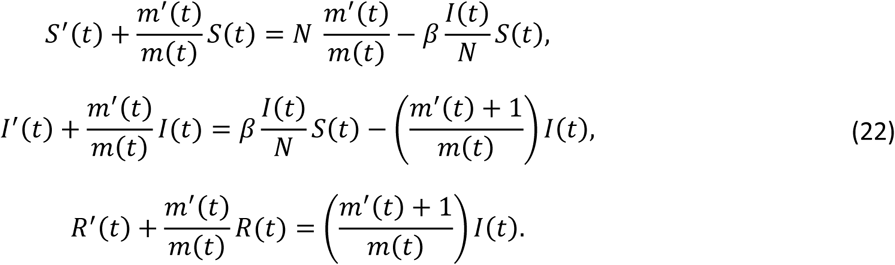

### 2.4 Equilibria, the basic reproductive number ℛ_0_, and the effective reproductive number ℛ_*e*_

The compartmental model (22) potentially have several equilibria. As is standard, (22) possess the standard disease free equilibrium:

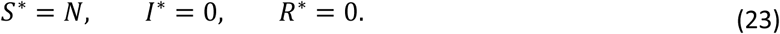

In addition, an equilibrium where the infection was exhausted, leaving susceptible individuals that escaped infection, and recovered individuals:

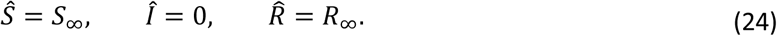

Turning our attention to the reproductive numbers of the disease, the basic reproductive number obtained directly from the survival function [28] is

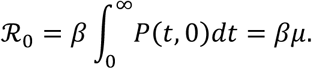

To estimate the basic reproductive number using the next-generation method, we have that 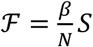 and 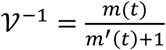. Note, the motivation for 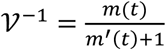 instead of 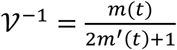 arises from 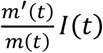 accounting for a change in average duration of infectivity, instead of the transfer of infected individuals to the recovered state. It follows that

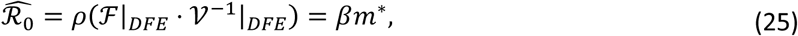

which is the spectral radius of the next generation matrix, ℱ|_*DEF*_ · 𝒱^−1^|_*DEF*_ and 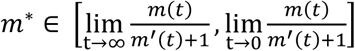.

Finally, we consider the effective reproductive number, in the face of time variability of *m*(*t*), *m*′(*t*), and *S*(*t*), to be

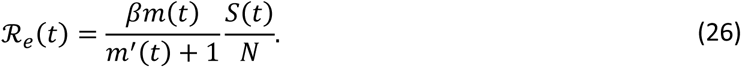

## 3. Special cases of the duration of infection distribution

We now consider the duration of infection distributions used in traditional differential equation compartmental models, namely the exponential distribution, and the Erlang distribution. In addition, we illustrate a compartmental model that accounts for any mean, standard deviation, skewness, and excess kurtosis, by assuming that the duration of infection is Pearson distributed.

### 3.1 The exponential distribution

If the duration of infection is exponentially distributed, then the associated survival function at the onset of the epidemic is *P*(*t, x*) = *e*^−*γ*(*t*−*x*)^. Under this assumption, we have that [29],

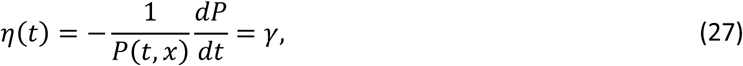

and

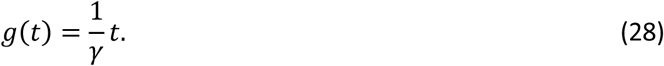

Solving (11) under the assumption of (28) yields,

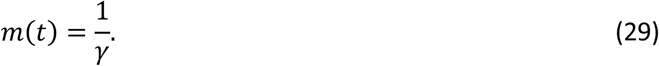

Substituting (29) into (22), we arrive at the traditional form of differential equation compartmental models:

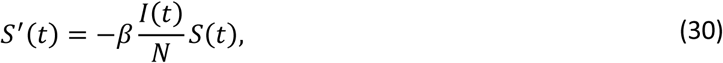

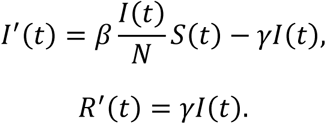

Finally, because 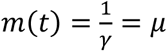, the basic reproductive numbers are

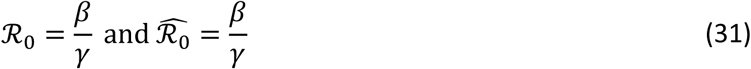

### 3.2 The Erlang distribution

If the duration of infection is Erlang distributed, then the survival function is

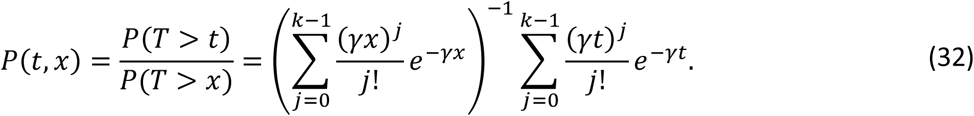

Note, this formulation of an Erlang distributed random variable differs from that used in the derivation of traditional compartmental models, as in general *P*(*t, x*) ≠ *P*(*t* − *x*).

From (9) and (32), it follows that

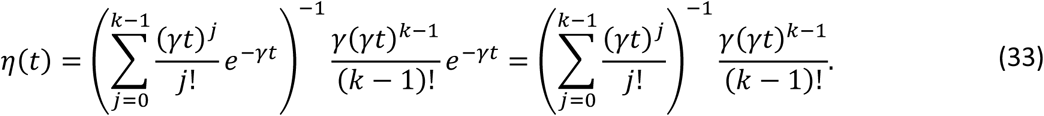

We also have that [29]

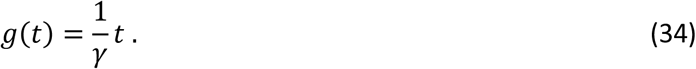

Substituting (34) in (11), we obtain the mean residual waiting-time:

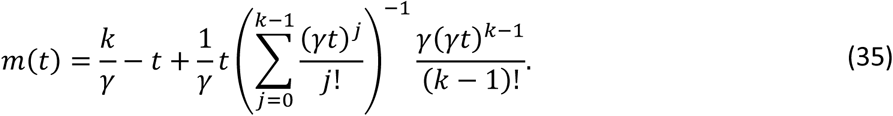

Using (35) and its derivative in system (22), we obtain a compartmental model comprised of three equations that features a duration of infection that is Erlang distributed, regardless of the value of *k*.

Subsequently by applying the ‘linear chain trick’ [9,31,32] (Supplemental Materials) to the system of three equations, we can illustrate the effects of a duration of infection that is Erlang distributed:

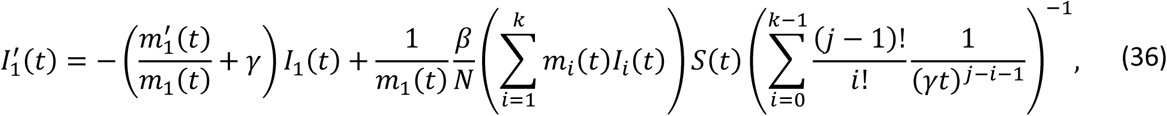

and

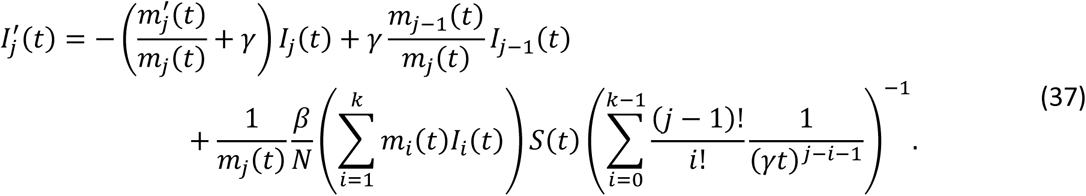

where *m*_*j*_(*t*) is the waiting-time of the *j*^*th*^ stage.

Finally, we consider the special case when individual stages feature identically constant waiting-times, which implies that

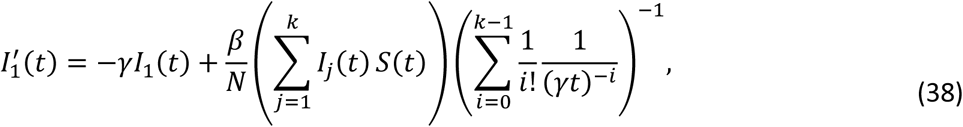

and

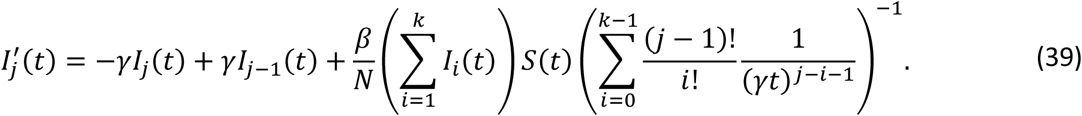

Through (32), (33) and (35), we reduce system (22) to a system of 3 differential equations based on an Erlang distributed duration of infection, or through the ‘linear chain trick’ a system of *k* + 2 differential equations (i.e. (46)-(47), *S*′ and *R*′). An important distinction between (38)-(39) and the traditional system of differential equations (6) with an Erlang distributed duration of infection is that new infections enter into any given infectious state in (38)-(39), based on a component of the duration of infection distribution, whereas (6) requires all new infections to enter the first stage of infection. Thereby, (38)-(39) present an approach that is likely to better conserve the variation in duration of infection, at least relative to its traditional compartmental model counterparts.

Finally, in regards to the basic reproductive numbers, we have that 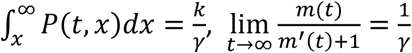 and 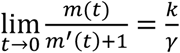,so

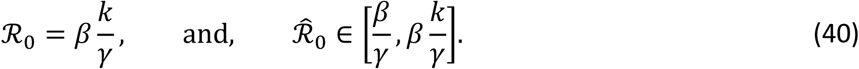

### 3.3. The Pearson distribution

If the duration of infection is Pearson distributed, then the associated survival function does not posses a general closed form. Thus, we define the Pearson distribution as

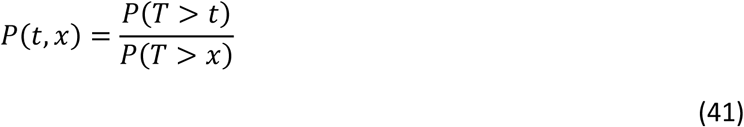

where

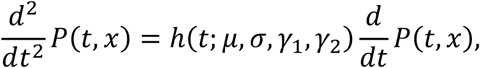

and

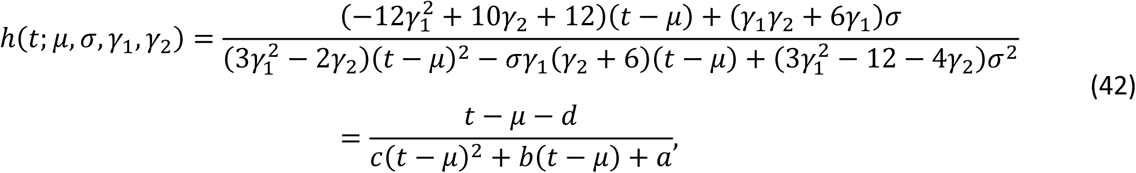

where 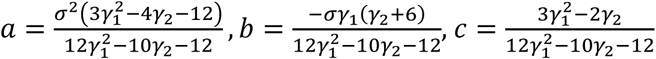 and 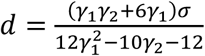.

Note, *μ, σ, γ*_1_, and *γ*_2_ are the mean, standard deviation, skewness, and excess kurtosis of the duration of infection [29,33,34].

To obtain the mean residual waiting-time in terms of the hazard function, we have that [29,35]:

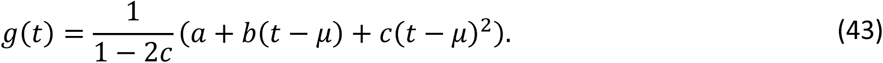

It follows from (18) that the mean residual waiting-time is:

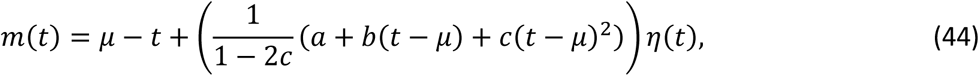

where *c* ≠ 1/2.

To ensure the mean residual waiting-time is finite we apply l’Hôpital’s rule to obtain

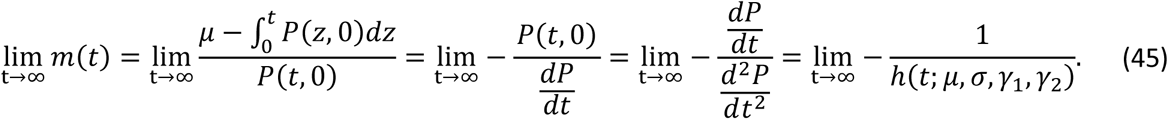

Thus, we require that 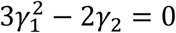 to force the terms involving *t*^2^ to drop out. Under this assumption, we have that

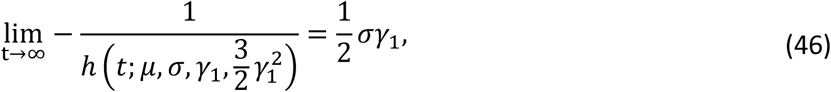

Given (46), the two formulations of the basic reproductive number are:

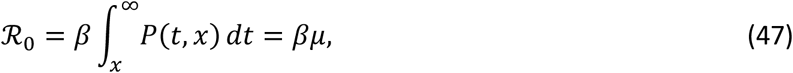

and

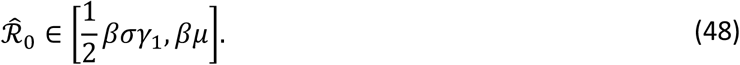

Note, the lower bound of (48) is consistent with the next-generation method estimate for the basic reproductive numbers (31) and (40), as the Erlang distribution has moments 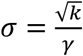 and 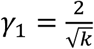, which implies 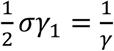.

## 4. Application of methodology

To illustrate the utility of our approach, we model measles outbreaks based on time series data of measles incidence in Iceland from 1924 to 1928 [36], taking into account classical data on the infectious period of measles [37].

We use sample moments of the mean, standard deviation, skewness, and kurtosis to estimate the infectious period distribution of measles, under the assumption that it is Pearson distributed, and compare the obtained novel compartmental model to a traditional compartmental model that features an infectious period that is Erlang distributed (Figure 1).

**Figure 1.**
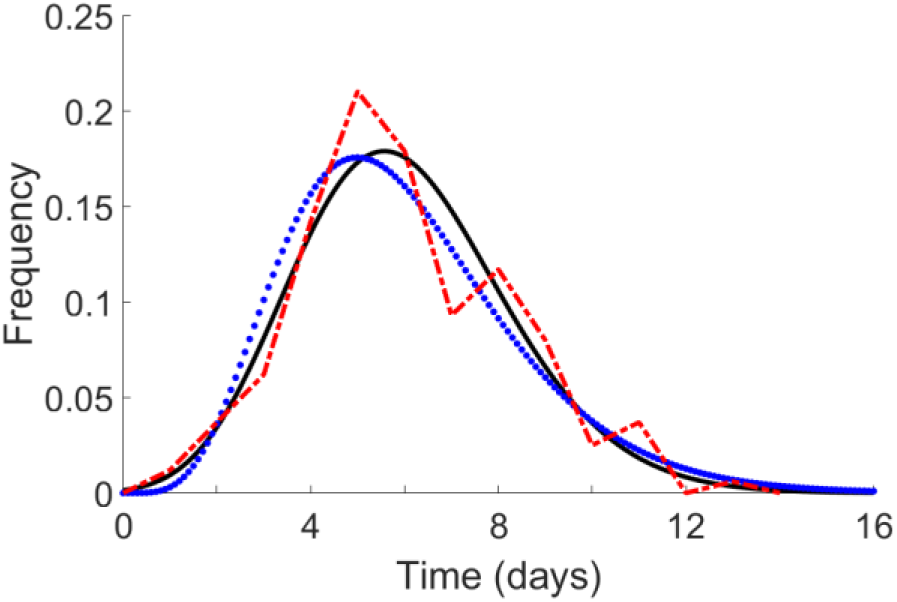
Measles infectious period. Infectious period distribution based on observed incidence data (red dash-dot line), fit Pearson distribution (black solid line) based on sample moments 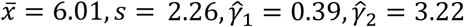, and fit Erlang distribution (blue dotted line) based a shape parameter of *n* = 6, and rate parameter of 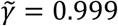.

For the novel compartmental model we determine the duration of infection distribution at fixed time points throughout the epidemic using (Figure 2):

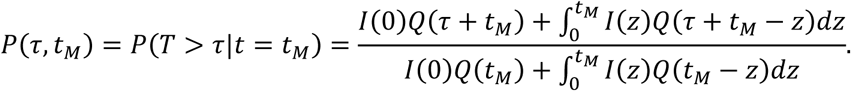

**Figure 2.**
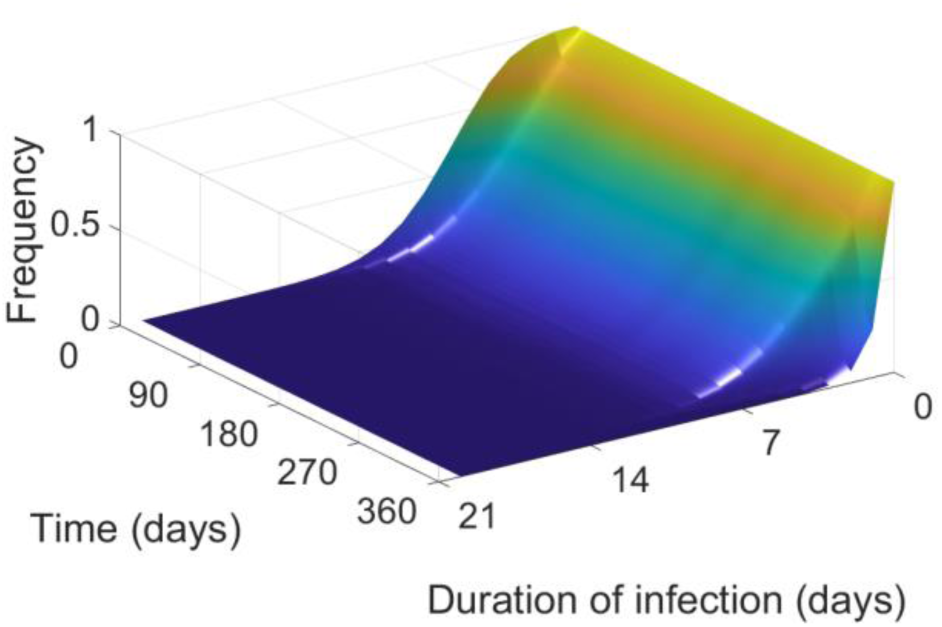
Survival function for individuals initially infected at given day throughout the epidemic.

Assuming that *P*(*τ, t*) is Pearson distributed with 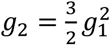, it follows that the mean, standard deviation, skewness, and kurtosis are 5.98, 4.15, 1.62, and 3.93 days, respectively. Given these moments, the hazard rate and mean residual waiting-time naturally follow (Figures 4). Finally fitting system (22) using a nonlinear least-squares procedure yields *β* ≈ 1.9061 (Figure 4).

**Figure 3.**
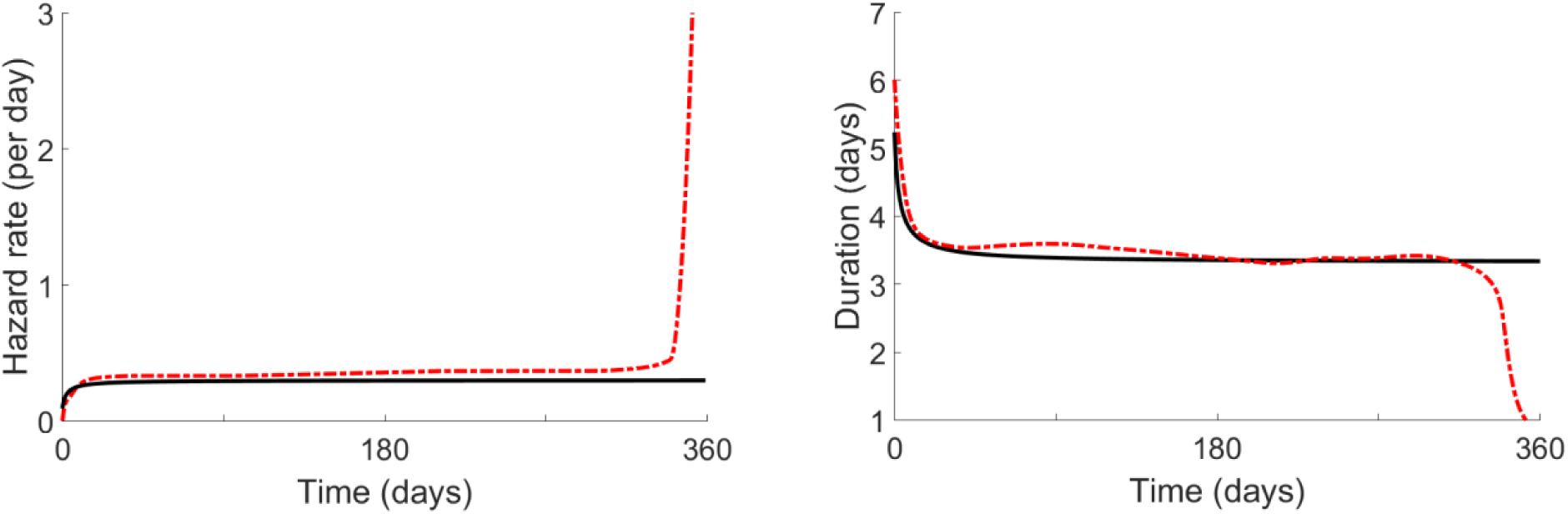
Hazard rate *η*(*t*) and mean residual waiting-time *m*(*t*).

**Figure 4.**
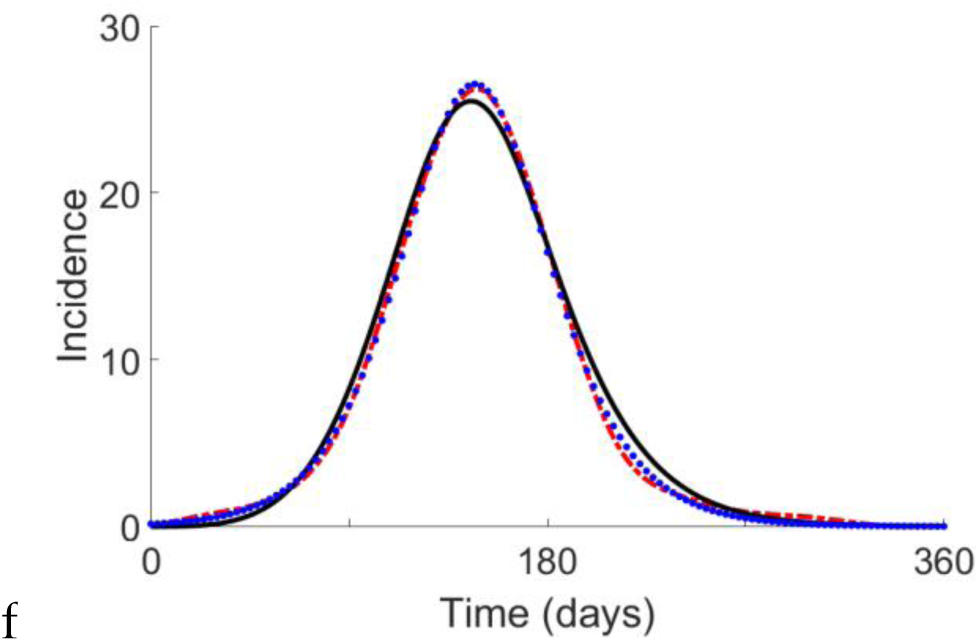
Predicted measles incidence in Reykjavik Iceland during 1924. Trajectories of incidence are based on interpolation of data (dash-dotted red line) [36], and simulated trajectory with novel compartmental model of SIR model (black solid line), and simulated trajectory with SI^k^R model (blue dotted line).

When *P*(*τ, t*) is Erlang distributed, we have that the mean and standard deviation are 5.98, and 2.44 days, respectively. Given these values, it follows that classical SIR model, with a duration of infection that is Erlang distributed, is

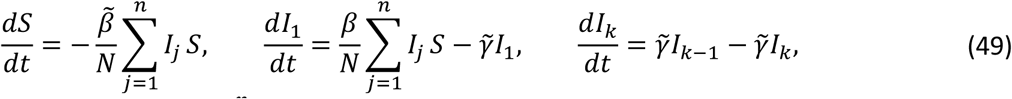

where *n* = 6, *μ* = 5.98 days, 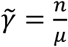, a non-linear least-squares fit (Figure 3) yields 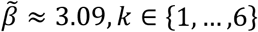, and the total population of Reykjavik (in 1924) is estimated to be *N* = 24134 [38].

Given the model parameters the possible range of basic reproductive number (47) of measles for the novel compartmental model (22) is given by

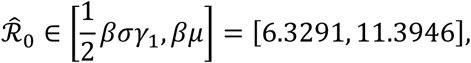

and for the traditional compartmental model (49),

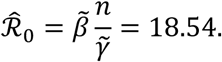

## 5. Discussion

In this work, we presented a class of novel differential equation compartmental models by modifying the classical assumptions that simplify nonlinear Volterra integral equations into systems of differential equations. To do this, we generalize the notion of the total infectivity of a disease, and the representation of its duration of infection. We illustrate the consistency of our class of novel models with traditional models for exponential and Erlang distributed durations of infections, present a new class of differential equation compartmental models based on a Pearson distributed duration of infection, and provide equilibria and basic reproductive numbers for our approach.

The requirement that the duration of infection follows an exponential distribution is often a source of weakness with regards to the biological validity of differential equation compartmental models. While the extension of such models through the linear chain trick [9,31,32] to a duration of infection that follows the Erlang distribution alleviates this weakness to some degree, it does so at the cost of inflating the size of the compartmental model, and thereby increasing the computational complexity of the system. Our new class of models avoids this inflation, while retaining the benefits of having a distribution of infection that follows an Erlang distribution. Thereby our new class of models offer an approach to reduce model complexity in an era when the complexity of compartmental models is ever increasing. Furthermore, if one generalizes the concept of a function to include the survival function of the Gamma distribution, our new class of models advantageously accounts for any Gamma distribution parameter values, including non-integer cases.

A main advantage of the new class of models is that they are ODE based. Therefore, like traditional compartmental models, they do not require specialist knowledge to use, and possess well-developed numerical methods for their simulation. Furthermore, the theoretical extensions of the traditional models to include alternative formulations of the force of infection, state-dependent recovery rates, in addition to the inclusion of additional disease compartments are easy to implement. In addition, many of the applications of the traditional models, such as the study of multi-strain dynamics, health benefit analysis, and cost effectiveness analysis, should naturally carry over without the need to reinvent the procedures of each analysis.

A potentially fruitful avenue for future applications of our new class of models is in the study of virulence and disease evolution. To elaborate, because our new class of model includes both the quantity of infected individuals and their duration of infection, it may serve as a better paradigm for investigating selective pressures that pathogens face, at least relative to traditional compartmental models. Similarly, our idea to track both the quantity of infected individuals along with their duration of infection could be adapted to study species competition, as a similar modification should provide stronger intuition on species fitness.

A surprising outcome of our work is the discovery that the lower bound provided by the next-generation method estimate of the basic reproductive number for the Pearson distributed example depends on standard deviation and skewness, instead of the mean. As standard deviation and skewness indicate the spread and lean of a distribution, it seems reasonable that their combination makes for a decent proxy for the location of the middle of a distribution. While this could be a consequence of assuming that the duration of infection follows the Pearson distribution, it also highlights a potentially new approach to bound the basic reproductive number for a disease directly from data.

The use of a duration of infection that is Pearson distributed may also open up an interesting avenue into bifurcation analysis. In particular, with modifications to incorporate demographic turnover or loss of immunity, and the use of the Pearson distribution, one could investigate if a relationship exists between the occurrence of periodic cycles and the first four moments of the duration of infection. Thereby, one may be able to gain intuition as to whether bifurcations are likely to occur simply by examining statistical moments and the diseases transmission rate. In addition, through such modified models, it may be possible to determine the existence (or non-existence) of hopf bifurcations by examining whether *m*(*t*) is periodic, as this seems like it would a requirement for such behavior in reality.

As our new class of models are based on the integral equation version of the Kermack and McKendrik model, it shares this model’s limitations. Namely, the assumptions of a sufficiently large and well-mixed population, the compartmentalization of diseases into distinct stages, and the transmission assumption of the law of mass-action. While these limitations may seem numerous, they do not impeded research on traditional models, and thereby should not inhibit the theoretical extension and application of the broader class of models proposed here.

The traditional assumptions that reduce the integral equation version of the Kermack and McKendrik model to a system of differential equations provides disease modellers with a rich source for mathematical and scientific discovery. Here, we proposed a generalization of these traditional assumptions to a biologically more accurate description of the total infectivity of a disease. By imposing these new assumptions, we provide a more descript picture of how a disease propagates throughout a population, while retaining the convenience and simplicity of differential equation compartmental models.

## Author’s contributions

SG conceptualized the project. CR and SG conducted formal analysis, wrote, reviewed, and edit the project manuscript.

